# Gene expression has more power for predicting *in vitro* cancer cell vulnerabilities than genomics

**DOI:** 10.1101/2020.02.21.959627

**Authors:** Joshua M. Dempster, John M. Krill-Burger, James M. McFarland, Allison Warren, Jesse S. Boehm, Francisca Vazquez, William C. Hahn, Todd R. Golub, Aviad Tsherniak

## Abstract

Achieving precision oncology requires accurate identification of targetable cancer vulnerabilities in patients. Generally, genomic features are regarded as the state-of-the-art method for stratifying patients for targeted therapies. In this work, we conduct the first rigorous comparison of DNA- and expression-based predictive models for viability across five datasets encompassing chemical and genetic perturbations. We find that expression consistently outperforms DNA for predicting vulnerabilities, including many currently stratified by canonical DNA markers. Contrary to their perception in the literature, the most accurate expression-based models depend on few features and are amenable to biological interpretation. This work points to the importance of exploring more comprehensive expression profiling in clinical settings.

## Introduction

Decades of work have confirmed cancer as a disease of the genome[1]. With this recognition has come the hope that the specific oncogenic mutations of cancer will confer equally specific vulnerabilities to therapeutic intervention. In the last decade, these vulnerabilities have been exhibited dramatically with compounds targeting known oncogenes such as B-RAF, EGFR, and ALK[2]. Driven by these early successes, the precision oncology paradigm has focused on sequencing patient tumors to determine the appropriate therapy[3]. However, genomically-indicated therapies have yet to benefit the majority of patients. Small proportions of patients have genomic indicators for a targeted therapy, and most that do respond for only a limited time, or in some cases do not respond at all[4].

Disappointing results with genome-based biomarkers have driven calls to look beyond the cancer genome to other possible indicators of cancer-specific vulnerabilities[5]. A major alternative for tumor characterization is the transcriptome. Although RNA is more difficult to interrogate in the clinic than DNA, studies have found that gene expression supplies the most significant predictive features for patient prognosis[6]. Previous work on predicting cell viability after compound treatment suggests that expression may be more powerful than DNA features for predicting drug response[7–9]. Additionally, several studies have combined individual gene expressions into gene set activation scores and found success in predicting compound response[10–12]. Despite these promising results, expression-based models are widely treated as undesirable either because they are considered trivial proxies for tissue-type[13] or because they are seen as uninterpretable[14]. However, Geeleher *et al.* demonstrated the potential clinical utility of expression data by showing that expression-based models trained on cell line drug response could predict clinical response to therapy[15]. More recently, the WINTHER trial of personalized cancer treatment included an arm to match patients to therapies based on their transcriptomic profile if they could not be matched using genomic alterations[16]. Including the transcriptomic arm increased the number of patients matched to therapies from 23% to 35% with no statistically significant difference in progression-free survival ratios between the genomic and transcriptomic arms.

These studies point to the need for a comprehensive comparison of expression and genomic molecular features as models of cancer vulnerability and a deeper interrogation of the interpretability of expression models. Here we present a study across five large datasets of cancer cell viability including both genetic and chemical perturbations. We find that RNA-Seq expression outperforms DNA-derived features in predicting cell viability response in nearly all cases, including many perturbations with known genomic biomarkers. The best results are typically driven by a small number of interpretable expression features. Our findings suggest that both existing and new cancer targets are frequently better identified using RNA-Seq gene expression than any combination of other cancer cell properties.

## Results

### Data and Methodology

We attempted to predict cell line viability in response to perturbations measured in five large *in vitro* cell viability studies, including CRISPR gene knockout (Cancer Dependency Map and Project Score[17]), arrayed oncology drug treatment experiments (GDSC[13]), pooled drug treatment experiments (PRISM Repurposing[18]), and RNAi knockdown studies (DEMETER2[19]). **(Fig. 1a-b**). Many perturbations do not produce differential viability effects across cell lines, indicating that no predictive model is likely to succeed. To assess performance specifically for cases where strong differential viability signal exists, we identified a subgroup of 100 perturbations in each dataset that appear to have the greatest biological signal based on their viability distributions across samples. We labeled these Strongly Selective Vulnerabilities (SSVs).

**Fig. 1:**
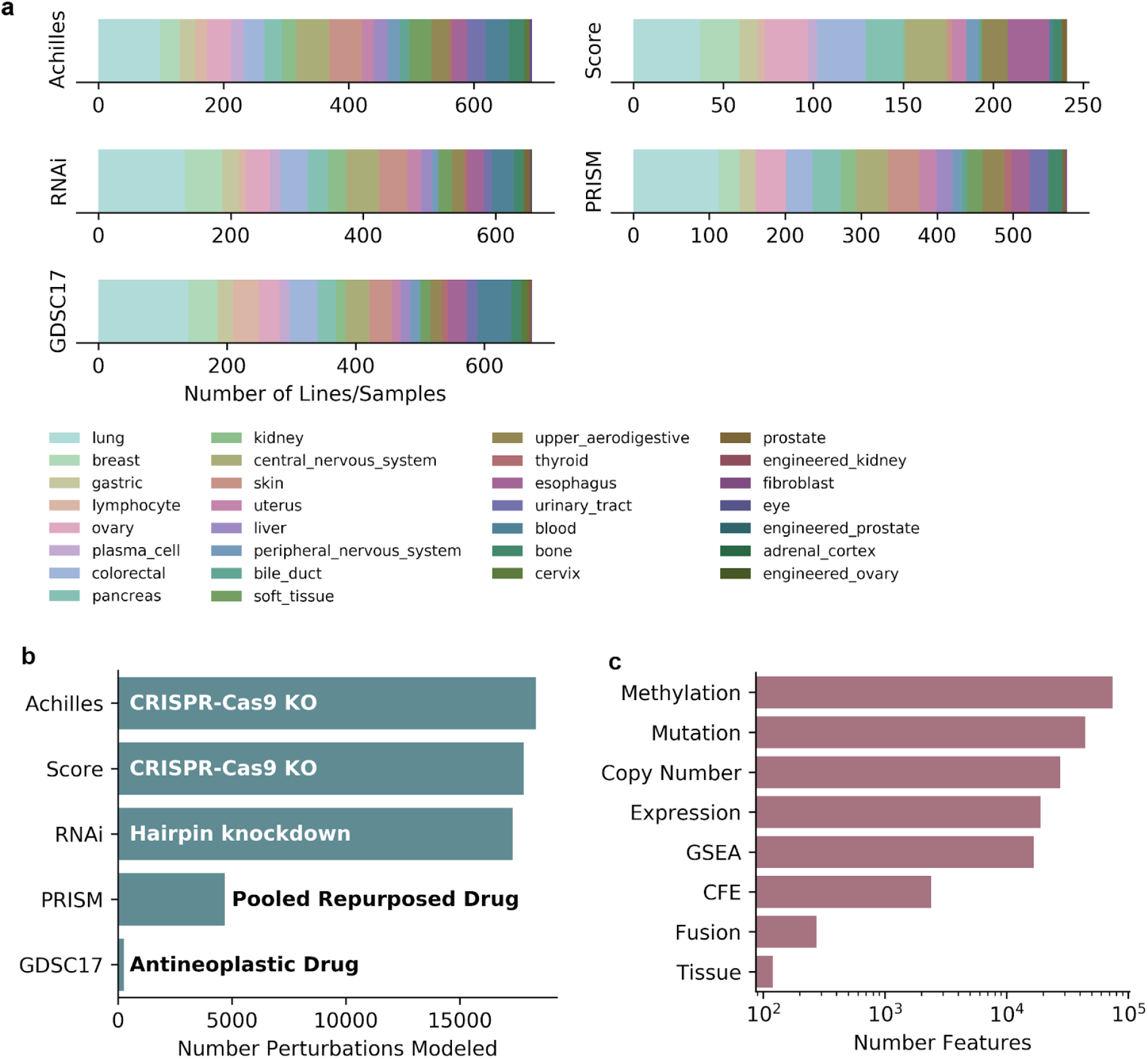
Overview of datasets analyzed. **a**. Breakdown of lineages covered by each dataset. **b.** Perturbations in each dataset where prediction was attempted. **c**. Number of CCLE features in each feature category. GSEA: Gene set enrichment analysis. CFE: Cancer Functional Events.

Molecular characterizations were drawn from the Cancer Dependency Map. They encompassed 181,951 total features, including mutation calls from whole-exome sequencing, RNA-Seq expression (protein-coding genes) at the single-gene and gene-set level, copy number alterations, reduced representation bisulfite sequencing (RBBS) methylation profiling, gene fusions, and tissue annotations (**Fig. 1c**). We refer to the union of all these measured properties as cell features. To see whether curation improved DNA performance, we adapted the Cancer Functional Events (CFEs) defined by Iorio *et al*. and introduced these as an additional genomics feature set[13], which we examine separately from the larger genomics feature sets in later analysis. After filtering for cell lines with characterization for mutation and RNA-seq expression data, the perturbation datasets spanned a set of 1,432 unique cell lines with 241-649 cell lines per dataset.

To evaluate the power of different cell properties for identifying the vulnerabilities of cancer cells, we trained a regression model individually for each perturbation. Our pipeline consisted of two components: a Pearson correlation filter to reduce the feature list to the top 1,000 features showing the strongest linear relationship to the viabilities, and a random forest model which produced the predictions using the selected 1,000 features (**Fig. 2a**). We found that not including a feature selection filter led to a slight drop in predictive performance due to the large number of irrelevant features included, along with a large increase in computation time. As in Iorio *et al*.[13] and Li *et al.[20]*, model predictions were scored using the Pearson correlation of the measured and out-of-sample predicted cell line viability measurements in each dataset, which we call the model’s Pearson correlation. A comparison of this random forest model with another commonly used machine learning model (elastic net) found that they produced similar performance on predicting SSV perturbations, but with a consistent edge for the random forest model (88.75% (355) of 500 SSVs better predicted by random forest; **Fig. 2 - figure supplement 1**).

**Fig. 2:**
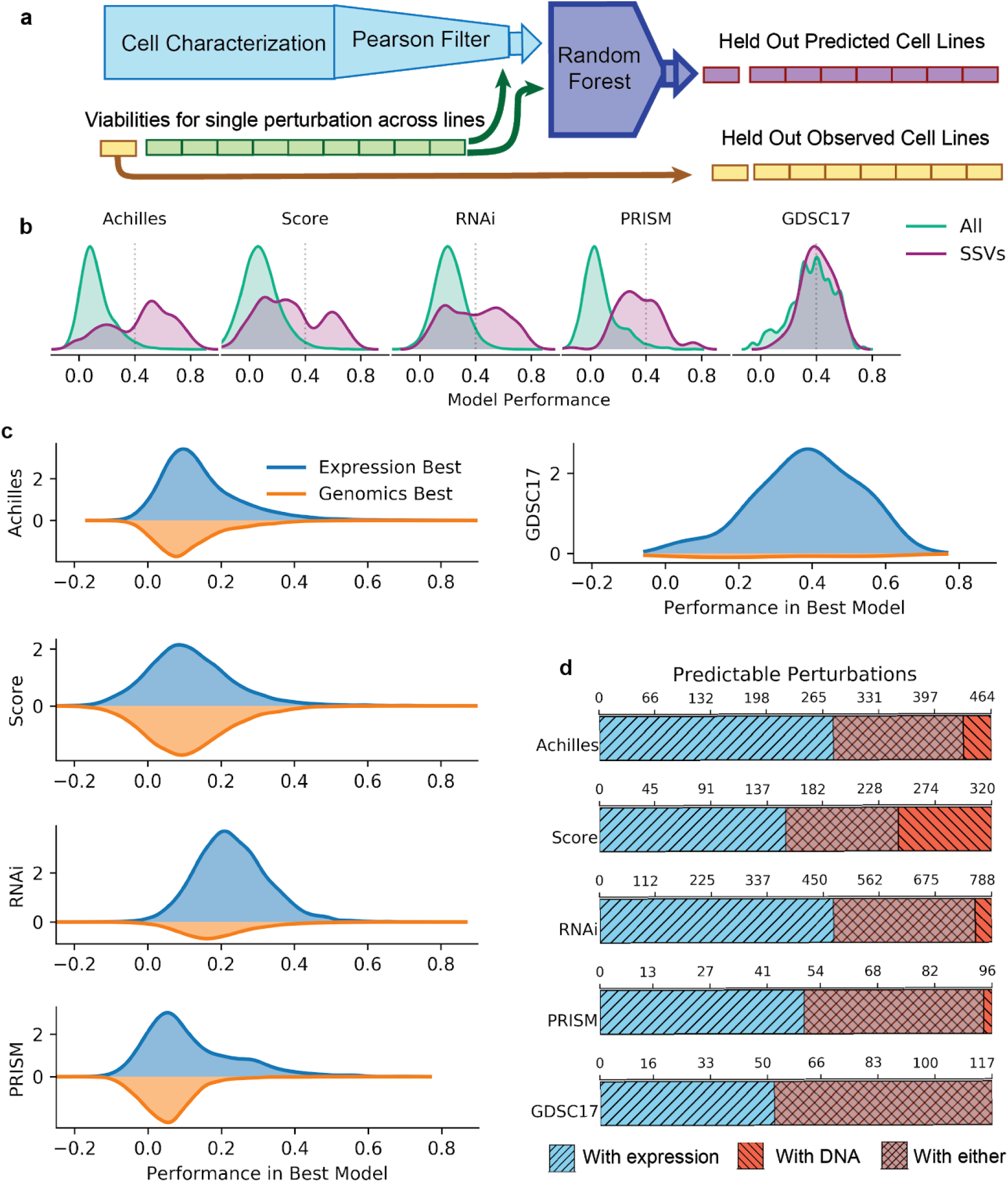
Prediction performance with DNA and expression features. **a.** Outline of the predictive algorithm used. Models are scored by the Pearson correlation of the concatenated out-of-sample predictions with observed values. **b.** Distribution of model predictive performance using all features for all perturbations (green) or SSVs (maroon). Dotted lines indicate the threshold to call a result minimally predictive (r=0.4). **c.** Performance distributions of the best of two models, one using all expression and one all genomic features. On top is shown the performance for cases where expression is best, on the bottom cases where genomics is best. **d**. Number of perturbations that can be predicted with a Pearson correlation of at least 0.4 with all expression or all DNA features. The cross-hatched overlap indicates the number of perturbations predictable at this threshold by either feature set.

### RNA-Seq Expression Features Outperform DNA-based Features

To establish baseline performance, we first examined predictive performance across all perturbations, using both DNA-based and RNA-based features in addition to annotations of primary tissue. Predictive performance varies strongly with the chosen dataset (**Fig. 2b**). Models predicting GDSC17 drug sensitivity data, which are focused on anti-cancer compounds, had the best overall performance (median Pearson correlation 0.387; 46.3% of perturbations predicted with a Pearson correlation greater than 0.4). Only 112 out of 4,686 compounds in the PRISM dataset had oncology as an annotated disease area, and unsurprisingly PRISM had the lowest overall Pearson correlations (median 0.048 and 2.2% above 0.4). As expected, prediction performance is improved and has less extreme variation across datasets among SSVs: 30% (Project Score) to 65% (Achilles) of SSVs were predicted with Pearson correlations above 0.4, with median Pearson correlations ranging from 0.265 (Project Score) to 0.490 (Achilles).

We then directly compared the performance of models supplied only expression features, which included RNA-Seq expression of individual genes and single-sample gene set enrichment analysis (GSEA) of MSigDB[21] gene sets against models supplied only DNA-based features: mutation, methylation, copy number, and gene fusions. Both feature sets included tissue annotations. Expression consistently outperformed genomics (**Fig 2c**). The benefit is especially pronounced when considering perturbations meeting some minimum threshold of predictability: using all expression features, a total of 1,641 perturbations across all datasets can be predicted with Pearson correlation at least 0.4, of which 1,001 (61%) cannot be so predicted by DNA features. Conversely, only 784 perturbations can be predicted with a Pearson correlation of at least 0.4 with DNA, and all but 144 (18.4%) can also be predicted by expression (**Fig. 2d**).

The selective killing profile of some perturbations is likely related to tissue-of-origin-specific rather than cancer-specific dependency. Accordingly, we compared performance in the Achilles CRISPR dataset specifically for the 278 genes with non-expression-based variants listed by OncoKB as “likely oncogenic” or “oncogenic”[22]. Even for the majority of these genes, expression features outperformed genomics for predicting viability after CRISPR knockout (**Fig. 2 - figure supplement 2**). The notable exceptions were BRAF, KRAS, NRAS (best predicted with their own mutation statuses), and SHOC2 (best predicted by NRAS mutation status). A similar pattern holds in the Project Score CRISPR and the RNAi gene dependency datasets, with most genes better predicted by expression but RAS genes and BRAF notably better predicted by their own mutation statuses.

Given that there is likely substantial redundancy between these DNA and RNA-based feature sets, we next asked what was the value of adding one feature set to the other for improving predictions. To address this question, we first took the twenty CRISPR perturbations best predicted with DNA features (**Fig. 2 - figure supplement 3a**) or expression (**Fig. 2 - figure supplement 3b**). We evaluated the change in performance when both feature sets were used for prediction. Adding expression improved the strongest DNA-based model performance by a median of 8.1% across datasets. In contrast, adding DNA features to the top expression models usually degraded performance by small amounts (due to the models overfitting slightly to noise in the datasets). Only RNAi showed a positive median change in performance when adding DNA features to expression (by 0.43%).

To further isolate each feature type’s contributions to the predictions, we next built models using each type of feature individually or in combination with single gene expression. All feature combinations included tissue annotations to ensure every model could identify the tissue of origin. Across all data sets, any feature combination that included single-gene expression produced equal or more minimally predictive (Pearson correlation greater than 0.4) models than any feature set that did not (**Fig. 3a**). For a given perturbation with at least one minimally predictive model using any feature set, the best model included expression in 76.3% (Project Score) to 99.2% (PRISM) of cases (**Fig. 3b**). These results make it clear that expression features are the key elements for successfully predicting viability responses to most perturbations, and are beneficial even when DNA features suffice for successful prediction.

**Fig. 3:**
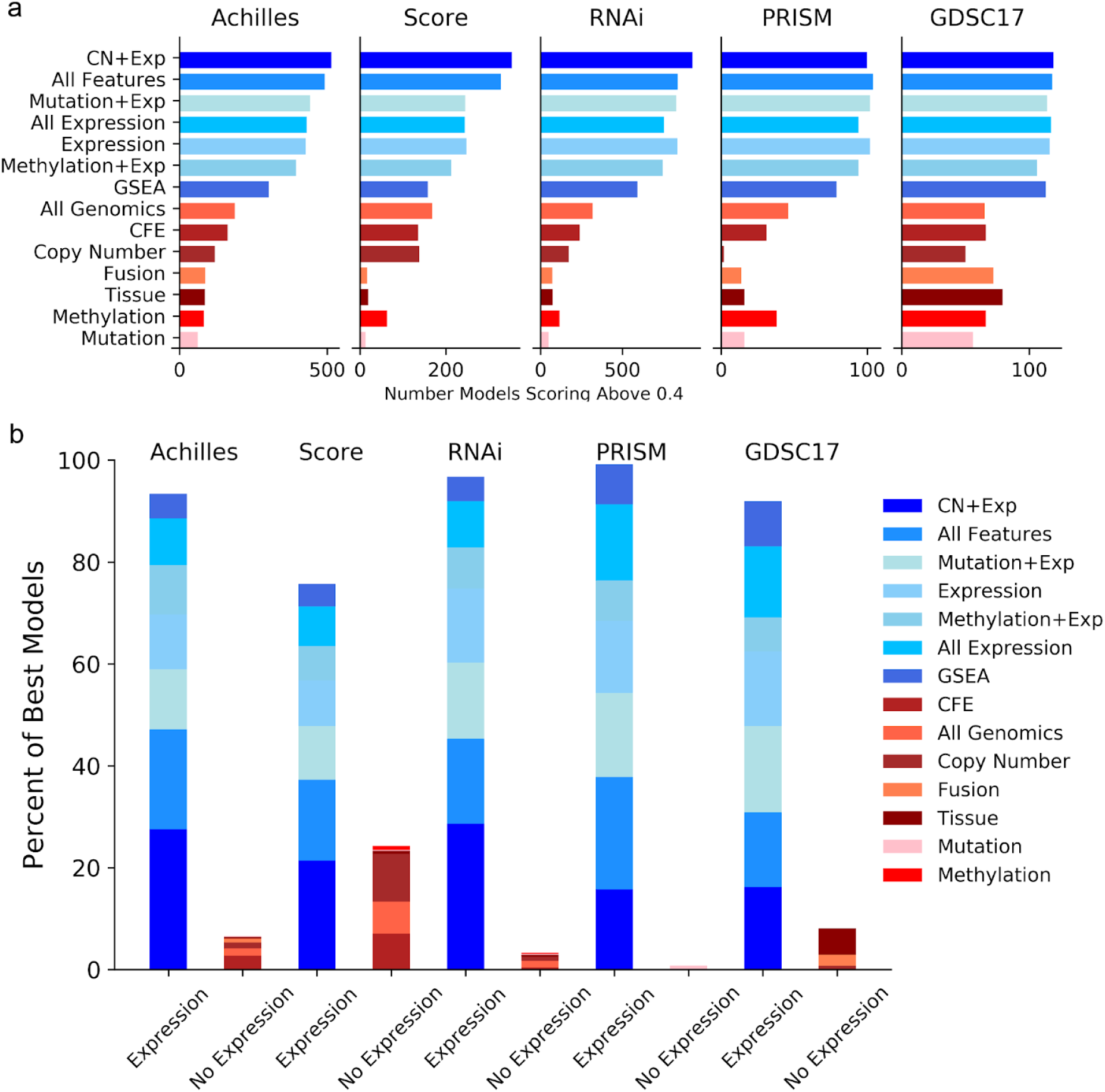
Performance of individual feature combinations. **a.** Number of predictive models achieving greater than 0.4 Pearson correlation with different feature combinations. CFE stands for Cancer Functional Events, a curated set of DNA features[13]. Expression includes only single-gene RNASeq profiles, while All Expression includes ssGSEA scores for MSigDB gene sets as well. **b.** For perturbations with at least one model with Pearson correlation above 0.4, percentage of models where each feature combination produced the best model.

### Specificity and Interpretability of Features

Models that utilize ‘diffuse’ sets of hundreds or thousands of features may be difficult to understand and act on. In contrast, models that use only a handful of features (sparse models) are generally easier to interpret, less vulnerable to overfitting, and potentially easier to measure in a clinical setting. We found that the most successful models trained with all features generally utilized a small number of features as indicated by the high importance they assigned to the top two features vs. the rest (**Fig. 4a**). Although predictive models using gene expression features are perceived as more diffuse and therefore less tractable than genomic models[14], we did not find this to be necessarily the case for either SSVs or minimally predictable perturbations. Sizeable fractions of both were dominated by one or more key features (features with more than 10% of total feature importance as assessed by sklearn’s Random Forest Regressor). Key features were often the expression of individual genes, even for models trained with all features including gene sets (**Fig. 4bc**). For the genetic dependency datasets only, we investigated how often key expression features were either the target gene’s own expression, that of a paralog[23], a strongly interacting protein[24], or of a complex co-member[25]. Combining these three measures, the median number of related genes per gene perturbation was only 29. Nonetheless, key single gene expressions were found to have a known relationship to the target in as much as half of gene dependencies (50.1% for Achilles, 29.3% for Project Score, and 40.7% for RNAi).

**Fig. 4:**
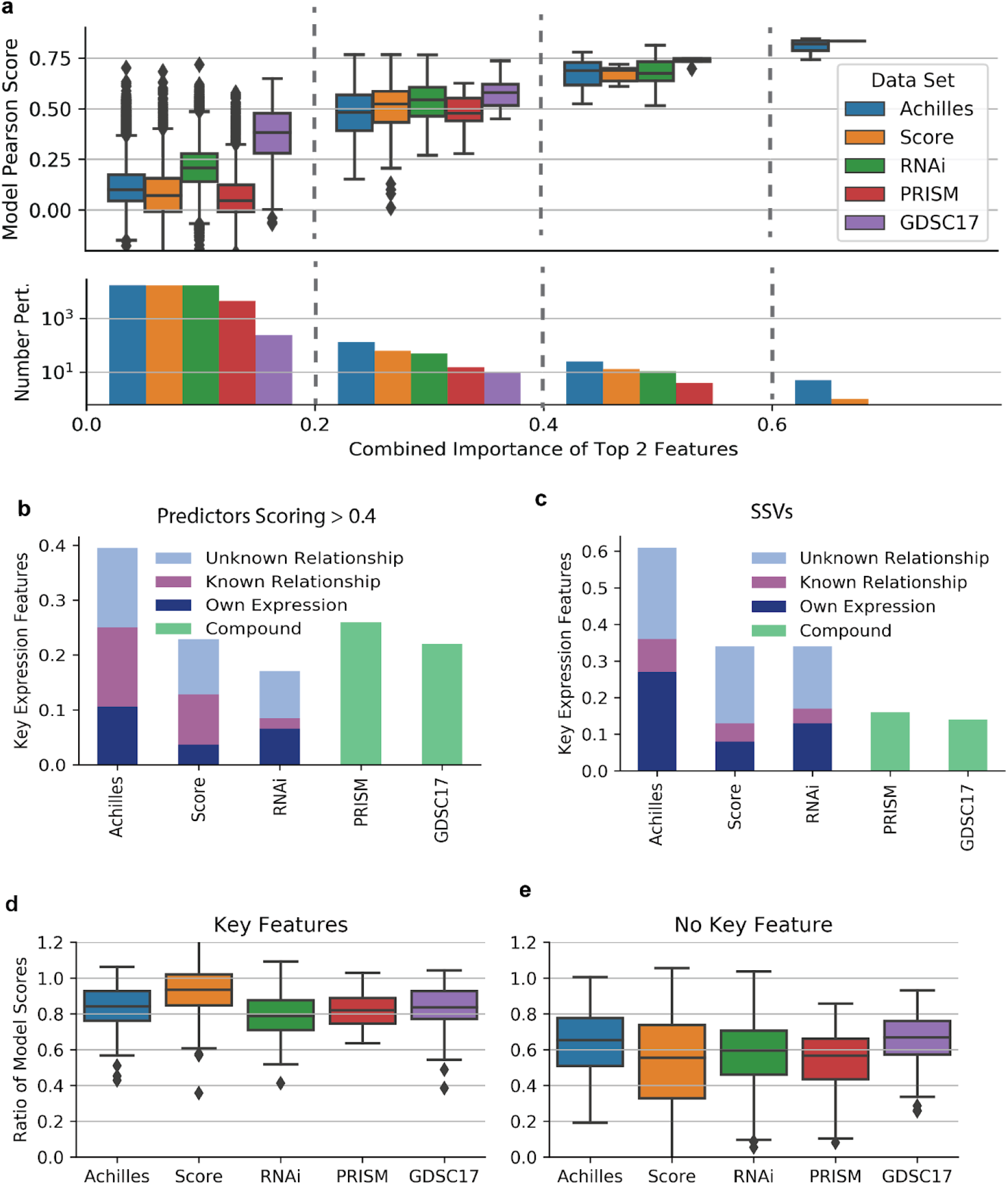
Sparsity of models. **a.** For models using all features, the combined importance of the top two features vs model performance. Perturbations are binned by top feature importance (dashed lines). **b**. For all-feature models with Pearson correlation above 0.4, fraction of models where the top feature is a single gene’s expression with at least 10% of total feature importance. For genetic perturbations, subcategories indicate the fraction known to be related to the target or the target itself. **c.** Similar results for SSV perturbations. **d.** The ratio of model accuracy using only the top feature (selected using random forest feature importance) or all features. Only models with Pearson correlation above 0.4 with a key feature are included. **e.** Similar for models with no key feature.

To confirm that apparently sparse models do not require many features to perform well, we examined how model performance varied with the number of features provided. For each training fold, we trained a model with the 1,000 features selected by Pearson correlation, ranked them by importance, and then trained using the top one, two, five, ten, or hundred features (**Fig. 4 - figure supplement 1**). We were specifically interested in whether models that appear to rely heavily on key features would retain their performance if they were supplied only a few features. We took models with at least a 0.4 Pearson correlation and at least one key feature with more than 10% of total importance among all features and examined performance with only the top feature (**Fig. 4d**). 62% of these single-feature models achieved at least 80% of their Pearson correlation using all features. Among perturbations predicted with greater than 0.4 Pearson correlation without a single key feature, only 14.0% could be similarly well-predicted with their top feature (**Fig. 4e**). With two features retained, 83.6% of models with key features achieved at least 80% of all feature performance (median performance ratio: 89.3%), while only 31.4% of models without a key feature achieved the same threshold (median performance ratio: 73.9%; **Fig. 4 - figure supplement 2**). We confirmed that these results were not a product of either the feature filtering strategy or the choice of random forest model by repeating this analysis with elastic net (EN) models for SSVs. We found that for perturbations where the random forest had identified a key feature, EN models using only the top feature achieved a median of 89.9% of their performance with all features (**Fig. 4 - figure supplement 2**). Taken together, these results suggest that the most predictable cancer vulnerabilities often require only one or two features for reasonable prediction.

Since single-gene expressions are highly structured and intercorrelated[26], some key expression features we identified are likely sufficient but not necessary for good prediction. We selected expression-only based models with at least one feature having greater than 10% of total importance and examined how performance changed when we removed that feature. Median predictive performance decreased by −1.8% (GDSC) to −26.7% (Project Score) (**Fig. 5**). The pattern of outcomes was very different for the different datasets, with nearly all PRISM perturbations showing some decline in performance while many GDSC perturbations appeared unaffected. In the two CRISPR datasets Achilles and Project Score, there was a clear trend where the best-predicted perturbations suffered the worst decrease in performance without their key features (Pearson correlation *p* = 3.98 × 10^−3^, 7.36 × 10^−7^ and *N* = 505, 290 respectively). All datasets showed a strong correlation between loss of performance and the importance of the top removed feature, with Pearson’s *R* ranging from −0.229 (RNAi) to −0.584 (Project Score) and the largest *p* value being 1.26 × 10^−5^ (RNAi, *N* = 66). Overall, 1,133 out of 1,260 (89.9%) expression models with a key feature lost performance when that feature was removed.

**Fig. 5:**
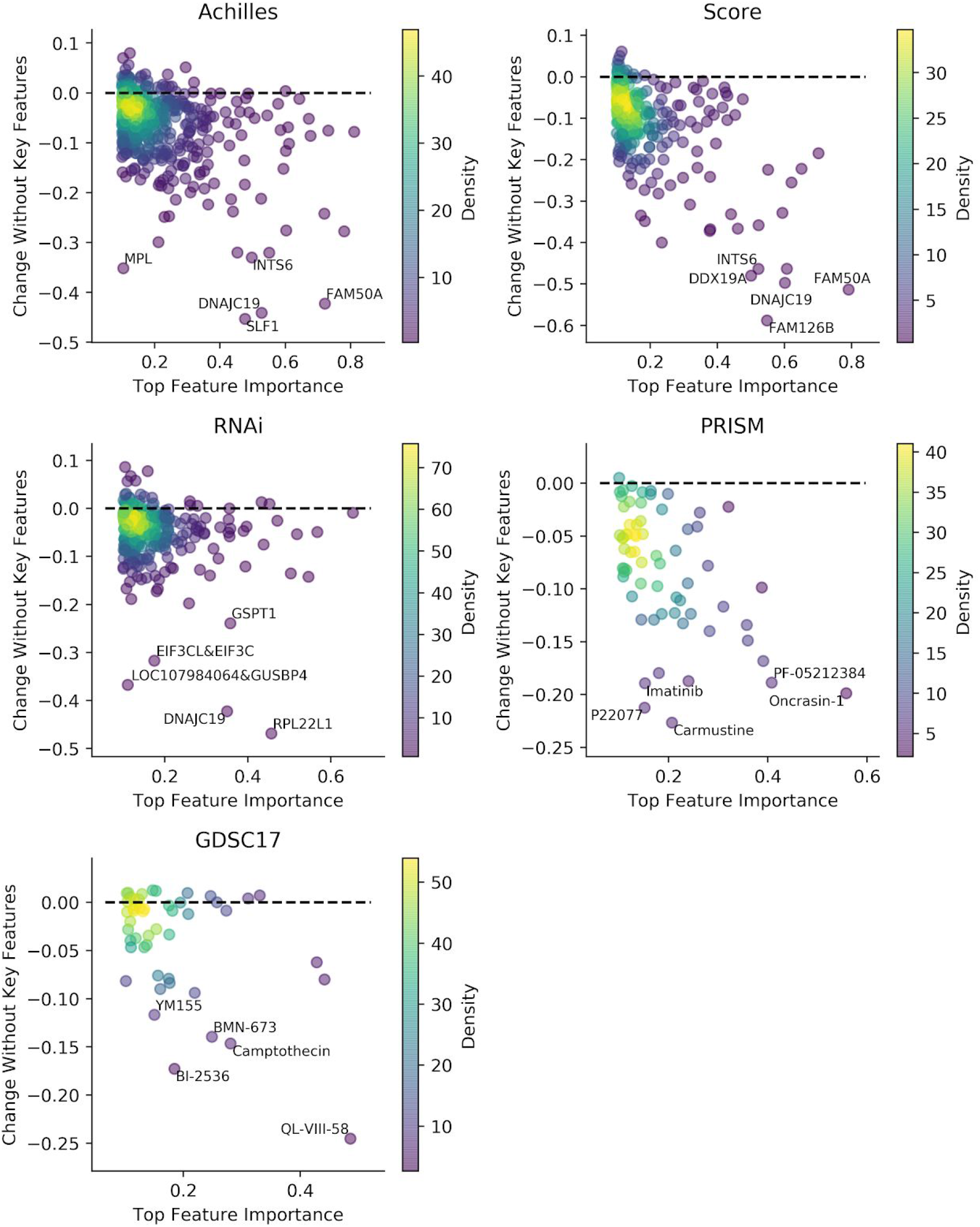
Effect of removing key expression features. For expression-only models with at least one key feature, we trained new models for the same perturbation that excluded all key features. We measured the change in the Pearson correlation (y axis) vs. the importance of the top feature in the original model.

By examining the models with the most severe performance loss after removing a top feature, we can learn which expression features are indispensable. In CRISPR, the most striking loss of performance generally occurred when a paralog’s expression was removed (e.g., FAM50A dependency with FAM50B gene expression removed). For the top five worst losses in GDSC17, the removed feature was either the expression of the multi-drug resistance gene ABCB1 (QL-VIII-58, BI-2536, YM155) or the expression of DNA helicase SLFN11 (camptothecin, BMN-673), whose expression is known to sensitize cell lines to DNA damaging agents[27]. In contrast, the most indispensable expression features for PRISM perturbations were more varied. For example, SULT1A1 expression was critical for predicting oncrasin-1 response, as would be expected from work by Huang *et al.*.28 Expression of GSTA1, which protects cells from oxidative stress, is essential for predicting sensitivity to the putative ubiquitin-specific protease inhibitor P22077.

### Interpreting models

To demonstrate how sparse models can be made interpretable, we trained individual regression trees using only features with importance greater than 0.08. This threshold and other tree hyperparameters were chosen to balance the tradeoff between simplicity and accuracy. In cases where the tree used only one feature, we illustrated the top feature’s relationship with the target. Some examples are discussed below.

TP53 loss-of-function mutation is one of the key molecular characteristics of cancer across lineage types. Surprisingly, for both Achilles and Project Score data, the top two markers of viability response to TP53 knockout did not include TP53 mutation status but the mRNA expression of EDA2R (a p53 transcriptional target), along with a gene set defined by the transcriptional response of skin cells to ionizing radiation[28] (**Fig. 6ab**). For RNAi-based suppression of TP53 the top model was a similar gene set defined by the response of primary blood mononuclear cells[29]. Both ionizing radiation gene sets include EDA2R. These results indicate that TP53 functional status is more cleanly inferred by these models from the transcription of its targets than directly from existing TP53 mutation calls. This may be due to failures in mutation calling or the complexity of possible causes of TP53 loss of function[30].

**Fig. 6:**
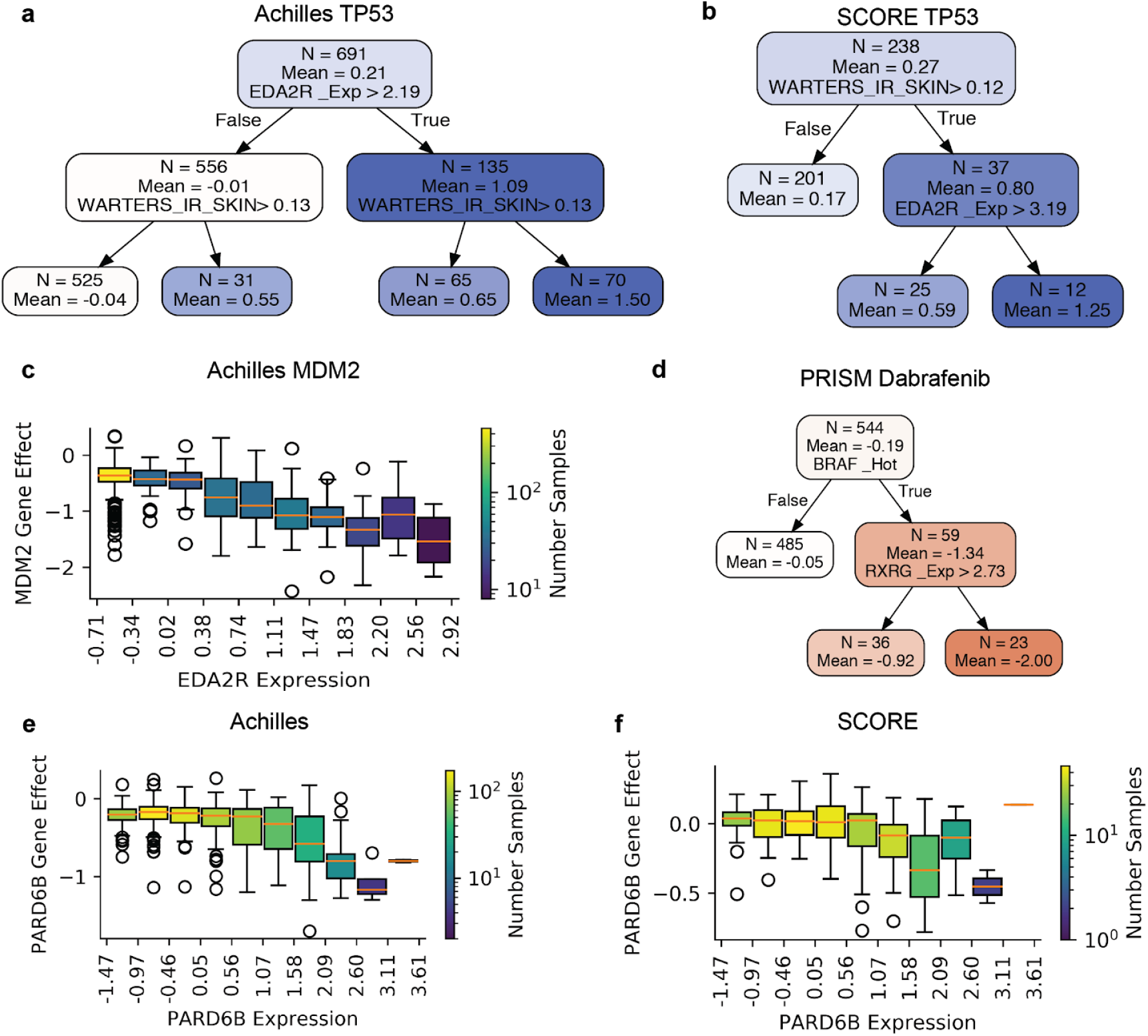
Interpretable models. **a-c**. Simple models for Achilles and Project Score viability post TP53 knockout and Achilles MDM2 knockout. **d**. Decision tree for PRISM dabrafenib response. **e-f**. Relationship between PARD6B knockout effect on viability and its own expression in Achilles and Project Score.

In cases where TP53 is functionally intact, cancers often depend on MDM2, a negative regulator of p53, to continue proliferation[31]. EDA2R expression was the top model of MDM2 dependency in Achilles (**Fig. 6c**) and was a top association in Project Score and RNAi data (**Table 1**). There was a clear gradation of response to either TP53 or MDM2 knockout according to the magnitude of EDA2R expression observed in all of these datasets (**Fig. 6 - figure supplement 1**). Response to the MDM2 inhibitor nutlin-3 also exhibited negative correlation with EDA2R expression in GDSC17 (**Fig. 6 - figure supplement 2**). The more potent MDM2 inhibitor idasanutlin was present only in the PRISM Repurposing dataset, where it also showed a strong relationship with EDA2R expression (**Table 1**). Given the significance of TP53 status in cancers, incorporating the expression of EDA2R and other TP53 targets for stratifying patients has considerable potential for clinical benefit.

**Table 1:** Relationship between selected perturbations and features. All *p*-values are calculated for the observed strength of Pearson correlation *ρ*.

| TP53 Pathway |  |  |  |  |  |  |
| --- | --- | --- | --- | --- | --- | --- |
| Dataset | Target | Feature | Context | $\rho$ | $p$ | $N$ |
| Achilles | TP53 | EDA2R Exp | All | 0.637 | $4.54 \times 10^{-80}$ | 693 |
| Score | TP53 | EDA2R Exp | All | 0.546 | $3.74 \times 10^{-20}$ | 241 |
| RNAi | TP53 | EDA2R Exp | All | 0.620 | $6.70 \times 10^{-71}$ | 655 |
| Achilles | MDM2 | EDA2R Exp | All | -0.661 | $1.61 \times 10^{-88}$ | 693 |
| Score | MDM2 | EDA2R Exp | All | -0.465 | $2.40 \times 10^{-14}$ | 241 |
| RNAi | MDM2 | EDA2R Exp | All | -0.616 | $9.56 \times 10^{-70}$ | 655 |
| GDSC17 | Nutlin-3 | EDA2R Exp | All | -0.495 | $1.34 \times 10^{-34}$ | 538 |
| PRISM | Nutlin-3 | EDA2R Exp | All | -0.358 | $7.31 \times 10^{-18}$ | 544 |
| PRISM | Idasanutlin | EDA2R Exp | All | -0.578 | $9.73 \times 10^{-52}$ | 566 |
| BRAF and BRAF Inhibitors |  |  |  |  |  |  |
| GDSC17 | Dabrafenib | RXRG Exp | BRAF Hot | -0.443 | $3.52 \times 10^{-4}$ | 61 |
| PRISM | Dabrafenib | RXRG Exp | BRAF Hot | -0.418 | $9.81 \times 10^{-4}$ | 59 |
| PRISM | Vemurafenib | RXRG Exp | BRAF Hot | -0.495 | $3.70 \times 10^{-5}$ | 63 |
| GDSC17 | Dabrafenib | RXRG Exp | Melanoma | -0.0182 | 0.922 | 61 |
| PRISM | Dabrafenib | RXRG Exp | Melanoma | -0.411 | 0.0103 | 38 |
| PRISM | Vemurafenib | RXRG Exp | Melanoma | -0.253 | 0.106 | 38 |
| Achilles | BRAF | RXRG Exp | Melanoma | -0.374 | 0.0113 | 45 |
| RNAi | BRAF | RXRG Exp | Melanoma | -0.386 | 0.0106 | 43 |
| PARDB6 |  |  |  |  |  |  |
| Achilles | PARD6B | PARD6B Exp | All | -0.444 | $7.73 \times 10^{-35}$ | 693 |
| Score | PARD6B | PARD6B Exp | All | -0.365 | $5.12 \times 10^{-9}$ | 241 |
| RNAi | PARD6B | PARD6B Exp | All | -0.309 | $2.78 \times 10^{-8}$ | 309 |
| Achilles | PARD6B | PARD6B Exp | OV+PANC | -0.544 | $2.80 \times 10^{-6}$ | 65 |
| Score | PARD6B | PARD6B Exp | OV+PANC | -0.434 | $2.89 \times 10^{-3}$ | 45 |
| RNAi | PARD6B | PARD6B Exp | OV+PANC | -0.559 | 0.249 | 6 |

Many canonical oncogenic gain-of-function mutations occur in the RAS-RAF pathway. We showed earlier that mutations in members of this pathway (BRAF, NRAS, HRAS, and KRAS) stood out in providing a signal of dependency that cannot be recovered from gene expression. However, it does not follow that gene expression has no value for predicting response to compounds targeting members of this pathway. For example, an interpretable model of the BRAF inhibitor dabrafenib in PRISM showed it is first stratified by BRAF mutation status, but the strength of response in BRAF-mutant lines is further stratified by expression of the nuclear factor Retinoid × Receptor Gamma (RXRG), which has been found to drive a neural crest stem cell state in melanoma[32,33] (**Fig. 6d, Fig. 6 - figure supplement 3**). The association of dabrafenib response with RXRG expression in BRAF-mutant lines was strong in both PRISM and GDSC17 (**Table 1**). A similar result was obtained in PRISM with the BRAF inhibitor vemurafenib (**Table 1**). As RXRG expression is enriched in melanoma, we checked whether these associations were still robust within the context of melanoma rather than BRAF hotspot mutations alone (**Table 1**). RXRG expression was significantly associated with increased dabrafenib sensitivity within melanoma in PRISM, but not in GDSC17. Additionally, vemurafenib sensitivity was negatively correlated with RXRG expression within PRISM melanoma lines, albeit not significantly (Pearson’s *r* = −0.253, *p* = 0.106). On the other hand, in both Achilles and RNAi, higher RXRG expression significantly predicted greater BRAF dependency within melanoma (**Table 1**). Project Score had only three annotated melanomas and was excluded. Overall, the evidence suggests that RXRG is a marker for increased BRAF dependency even within melanomas, but more screens in melanoma lines are needed to make a definitive conclusion.

In addition to stratifying patients for existing therapies, gene expression features could be critical for identifying sensitive populations for many new selective dependencies. Among these Par-6 family cell polarity regulator beta (PARD6B) which is predicted by its own expression. Cell lines with high expression show increased dependency in Achilles, Project Score, and RNAi (**Table 1, Fig. 6gh**). The effect is particularly pronounced within the ovarian and pancreatic cell lines: all three datasets increase correlation when restricting cell lines to these lineages (**Table 1, Fig. 6 - figure supplement 4**).

## Discussion

Measurement of gene expression is a potent and still under-utilized method for identifying vulnerabilities, with superior performance over genomic features in both genetic and compound response prediction. The advantage of expression-based features over DNA-based features held consistently across five high-throughput experimental platforms using three different perturbation technologies; multiple different subsets of perturbations, including cancer genes with known genomic indications; and multiple combinations of feature sets. Critically, expression-based models need not be black boxes. Many high-performing expression-based models we identified leveraged a small number (i.e., 1 or 2) of individual gene expression values and could be converted to easily interpretable models suitable for stratifying patients.

Iorio *et al.* also found that single-gene expression outperformed DNA-based features, but attributed this to the fact that RNA encodes tissue of origin[13]. When they attempted prediction within lineages using DNA-based Cancer Functional Events (CFEs) they found greater performance than with expression features. However, Rydenfel *et al*. found that expression and proteomics were more predictive than DNA even within tumor types[34]. We believe the discrepancy between these two studies can be explained by the strong curation applied by Iorio *et al.* to their DNA features. They identified a total of 1,063 DNA-based CFEs observed in at least one cell line[13]. This curation is quite successful in enriching for established drug targets: for example, 20.4% (54) of the compounds assayed in GDSC17 target a gene represented among the 472 cancer genes identified by Iorio *et al*. It is therefore reasonable that curated, binarized DNA features outperform uncurated, continuous, genome-wide expression in the context of very small sample sizes and compounds specifically developed against known genomic drivers. In such cases, it is more likely for a model to identify a genuine relationship if the number of irrelevant features is reduced through curation. Developing a similar curation strategy for expression features is expected to be fruitful for finding models in the context of limited sample sizes.

Some studies have suggested that the expression of gene sets, such as pathway activations[11] or inferred transcription factor activity[35,36], are more robust and interpretable predictors than expression of individual genes. We found that single genes’ expression data produced notably better results than gene set enrichment scores overall, despite having many more presumably irrelevant features. This result may be contingent on our particular choice of gene set enrichment algorithm (ssGSEA) and the gene sets considered (MSigDB). However, our study of the necessity of individual gene features found many cases in which missing a single key gene expression significantly undermined performance. Furthermore, predictive models for many vulnerabilities exhibit relationships we would expect to be specific to a single gene’s expression, such as when loss of a paralog’s expression predicts dependency or expression of the drug transporter ABCB1 predicts insensitivity to multiple agents. Overall, our study suggests at a minimum that single gene expression profiles have a value for predicting many perturbations that is not easily captured by simple measures of gene-set level expression.

We found in both PRISM and GDSC that elevated RXRG is a highly significant model of sensitivity to BRAF inhibition even within the setting of BRAF-mutated cancers. A study by Rambow *et al.* found using single-cell RNA-Seq that RXRG expression increases in melanoma BRAF^V600E/K^ patient-derived xenografts (PDXs) in response to anti-BRAF therapy and drives them into a neural crest stem cell state.[32] However, the authors identified this response as a key resistance mechanism to BRAF inhibitors, in contrast to results seen in PRISM, Achilles, and RNAi within melanoma. It may be that RXRG upregulation is a broadly protective response for stressed melanoma cells. In that case, constitutively high RXRG expression could indicate existing stress that leaves the cells with less reserve for additional challenges such as anti-BRAF therapy. Alternatively, the difference between studies may relate to the difference between PDXs and established cell lines or the difference between the *in vivo* and *in vitro* settings.

Although RNA-Seq is not yet a standard clinical diagnostic tool, it is rapidly advancing towards maturity. Widespread adoption of this technology has positive implications for assigning existing therapies to patients and finding new targets. However, this work suggests that a full RNA-Seq profile of tumors is not necessary to gain substantial benefits for precision therapy. Since many new or established vulnerabilities can be identified with just one or two expression features, cheaper technologies such as qPCR or FISH imaging against identified biomarkers would still unlock considerable benefits.

## Methods

### Data and Code Availability

The input matrices for the predictive models and all results needed to reconstruct the figures will be available on Figshare on publication. The code for the filtered random forest model and a jupyter notebook that generated the figure panels will be made available on github on publication.

### Preprocessing of Perturbation Data

CRISPR data were taken unaltered from the gene_effect file in avana_public_19Q2. RNAi data were taken from gene_effect in the combined RNAi dataset[37]. PRISM data were taken from Corsello *et al*.[18] GDSC17 data were downloaded using the GDSC data repository and processed using the gdscIC50 repository[38] following the “gdsc_17” vignette until data were normalized by dose such that negative controls were 0 and positive controls 100. For each compound, a single dose was chosen that had the greatest variance over cell lines for that compound, subject to the requirement that at least 100 cell lines have scores for that dosage in that compound. Cell lines were mapped to CCLE names using their COSMIC IDs. Project Score data were taken from CERES-processed gene effect scores[39].

### Preprocessing of Cell Features

Cell features were taken from the Cancer Dependency Map 19Q2 dataset.[40] Cell lines that did not have all four of RNA-Seq expression, mutation, copy number, and disease annotations were dropped. Categorical features were expanded in one-hot encodings. Features with zero variance in the remaining cell lines were dropped. Continuous features were Z-scored individually. Finally, all remaining missing values were filled with 0. Thus, the number of cell lines used for evaluating perturbations is the same regardless of the combination of features chosen for training.

Mutation data were divided into three cell-by-gene binary matrices with the following logic:

- Damaging: True if the gene has a deleterious mutation in the cell line.
- Hotspot: True if the gene has a non-deleterious mutation in a TCGA or COSMIC hotspot.
- Other: True if the gene has a mutation not falling in the other categories

For each MSigDB gene set, the R method GSVA::gsva was used on TPM RNA-Seq expression for CCLE lines released as of 19Q1.

To develop a curated set of genomic Cancer Functional Events similar to those identified by Iorio *et al.[13]*, we did the following:

- We took the 461 genes identified by Iorio et al. as high confidence cancer driver genes (level A, B, or C) and filtered both damaging and other matrices to only these genes.
- We took the recurrent copy number alterations (CNA) identified by Iorio et. al. and estimated the mean copy number of the corresponding segment in each cell line, using the WES-prioritized segmented copy number file from CCLE. We then produced a line-by-segment continuous matrix with the mean copy number values.
- We took the informative CpG islands identified by Iorio *et al.[13]*and produced a line-by-island matrix with the RBBS methylation values

CFE features with no results for any cell lines in CCLE were dropped. Cell lines with any missing values in any of the remaining features were dropped when training models using CFEs only.

### Identifying Interesting Perturbations

For CRISPR and RNAi data, we generated probabilities of dependency for each gene in each cell line using the methodology described in Dempster *et al*.[41] Only genes with at least five lines with greater than 0.5 probability of dependency and five lines with less than 0.5 probability of dependency were retained for training.

We identified strongly selective vulnerabilities (SSVs) in CRISPR, RNAi, and PRISM using the NormLRT score method developed by McDonald *et al.* [13,42] This test compares the quality of the fit of the perturbation’s distribution with a normal and skewed-t distribution, with higher scores indicating less normally-distributed data. The 100 perturbations with the highest scores in each dataset were labeled SSVs for the purposes of the following analyses. As noted above, GDSC data is scaled and clipped to the interval [0, 100], and therefore tests for normality on these data are inappropriate. Instead, we selected the 100 compounds with the greatest variance as SSVs for this dataset.

### Standard model Pipeline

For each perturbation evaluated, cell lines with missing perturbation scores or no feature values were removed. We then selected features belonging to the feature types being evaluated and split cell lines into ten equal folds, choosing one fold to leave out for evaluation and taking the rest for training.

Next, if the number of features was greater than 1,000, we filtered for the 1,000 features with highest absolute Pearson correlation to the perturbation scores in the training cell lines. We trained sklearn random forests with max depth eight, 100 trees, and a minimum of five cell lines per leaf using the training data and predicted perturbation viabilities for the held-out fold.

We repeated this process again, choosing a different fold to hold out evaluation, until we had out-of-sample predictions for all cell lines. We scored the model using the correlation of these out-of-sample predictions to the original perturbation scores.

For genes found to be predicted with Pearson correlation above 0.4, we repeated the training process, but this time additionally selected the top 1, 2, 5, 10, and 100 most important features per fold and trained models on just those features, saving their predictions on the held-out fold.

### Elastic Net model Pipeline

For each dataset, we adjusted all post-perturbation viabilities and all features to have mean 0 and variance 1. To reduce the cost of training the models, we chose fixed hyperparameters for each dataset in the following way: we took a random set of 20 SSVs and used the python implementation of glmnet to find the optimal value of lambda (the overall strength of regularization) with the ratio between L1 and L2 priors held fixed at 0.5, and nine specified lambda values spaced logarithmically in the interval [0.05, 50]. The optimal value of lambda was highly bimodal, with SSVs successfully predicted tending to small lambda values while those that could not be predicted using the maximum available value of lambda. We chose the mode of the smaller lambda values for training the dataset. This was 0.5 for Project Score data and 0.2 for all other datasets.

Training and scoring models proceeded as with the standard model pipeline, except that only SSVs were used and the magnitude of the coefficients was used to rank features instead of importance.

### Feature Importance

To assess feature importance for a perturbation, we performed Pearson feature filtering and trained a random forest using all available cell lines with feature and perturbation data. Feature importance was taken from the resulting model using sklearn’s default method, which measures gini importance. The resulting importance is normalized so that the sum of the importance of all features (of the thousand retained after Pearson filtering) is 1. Features with at least 0.1 importance were considered key features.

### Gene Relationships

Gene relationships were taken from the union of four relationships: the gene itself, its complex comembers, its protein interactors, and its paralogs. Complexes were taken from the CORUM file coreComplexes_03.09.2018_Corum_3.0.txt[25]. Protein-protein interactions were taken from InWeb_IM[24]. Paralogs were found with Ensembl Compara, and filtered for genes that underwent a duplication rather than speciation event[23].

### OncoKB Oncogenes

A table of all cancer gene variants and their significance was downloaded from oncokb.org on May 9th, 2018[22]. This table was filtered for genes with variants that were oncogenic or likely oncogenic and for which there was at least one non-expression based indication.

### Interpretable Models

We formed interpretable models of selected perturbations using single decision trees. We took the top three features identified by random forest. For each of the three chosen features, if it was continuous, we trained a depth-1 sklearn RegressionTree to predict the perturbation and used the resulting optimal split to binarize the feature. We then trained an sklearn RegressionTree with max depth 4 and minimum samples per leaf 5, and the results visualized as in **Fig. 6**. The trees were additionally regularized to require a minimum impurity decrease per split of 0.02 * *S*. *S* is a standard deviation found by taking all successfully predicted perturbations in the given dataset with a key feature, ravelling all their viability measurements into a single long vector, and calculating the standard deviation.

## Data and Code Availability

All supplementary data needed to reproduce these results, a python package for training and evaluating the described elastic net and filtered random forest models with various feature subsets, and an interactive jupyter notebook that generates all plots shown in this manuscript will be made available with publication.

## Author Contributions

J.D., A.T., and T.R.G. conceived the study. J.D. collected the data, trained models, and performed the analyses. J. K.-B. created the norm LRT calculation pipeline. A.W. performed ssGSEA calculations to generate gene set enrichment features. J.M., T.R.G., J.S.B., F.V., W.C.H. and A.T. supervised the study.

## Acknowledgments

The authors would like to thank Francesco Iorio and Emanuel Gonçalves for helpful commentary and suggestions.

## Competing Interests

T.R.G. is a paid consultant to GlaxoSmithKiline, is a co-founder of Sherlock Biosciences and FORMA Therapeutics, and receives sponsored research funding from BayerHealthCare, Calico Life Sciences and Novo Ventures. A.T. is a consultant for Tango Therapeutics. W.C.H. is a consultant for ThermoFisher, Solasta, MPM Capital, iTeos, Frontier Medicines and Paraxel and is a Scientific Founder and serves on the Scientific Advisory Board (SAB) for KSQ Therapeutics. F.V. receives funding from Novo Ventures. All other authors declare no competing interests. All authors were partially funded by the Cancer Dependency Map Consortium, but no consortium member was involved in or influenced this study.

---

**Fig. 2 - figure supplement 1:**
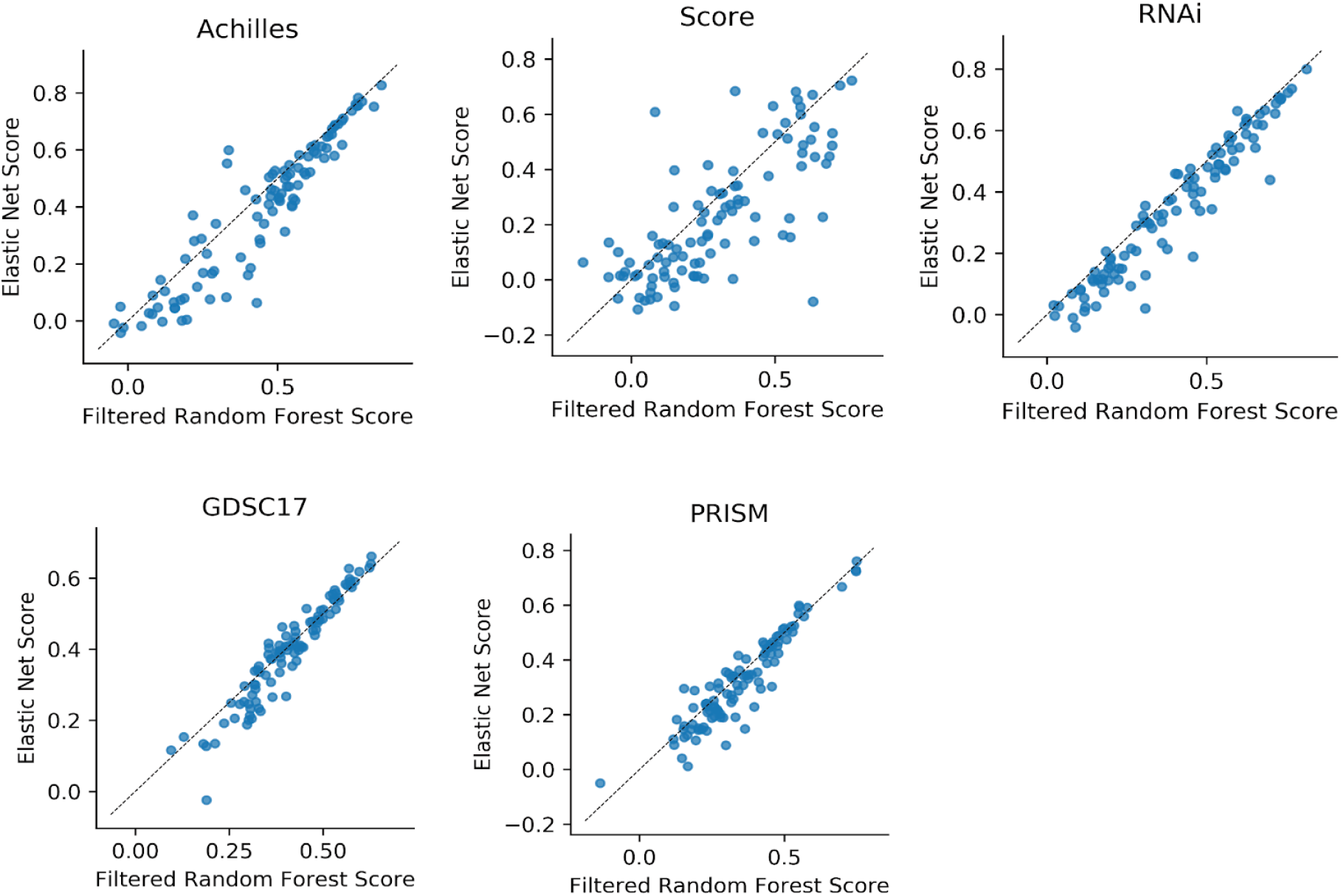
Comparison of random forest with feature preselection and elastic net. For each dataset, the Pearson score for SSV perturbations.

**Fig. 2 - figure supplement 2:**
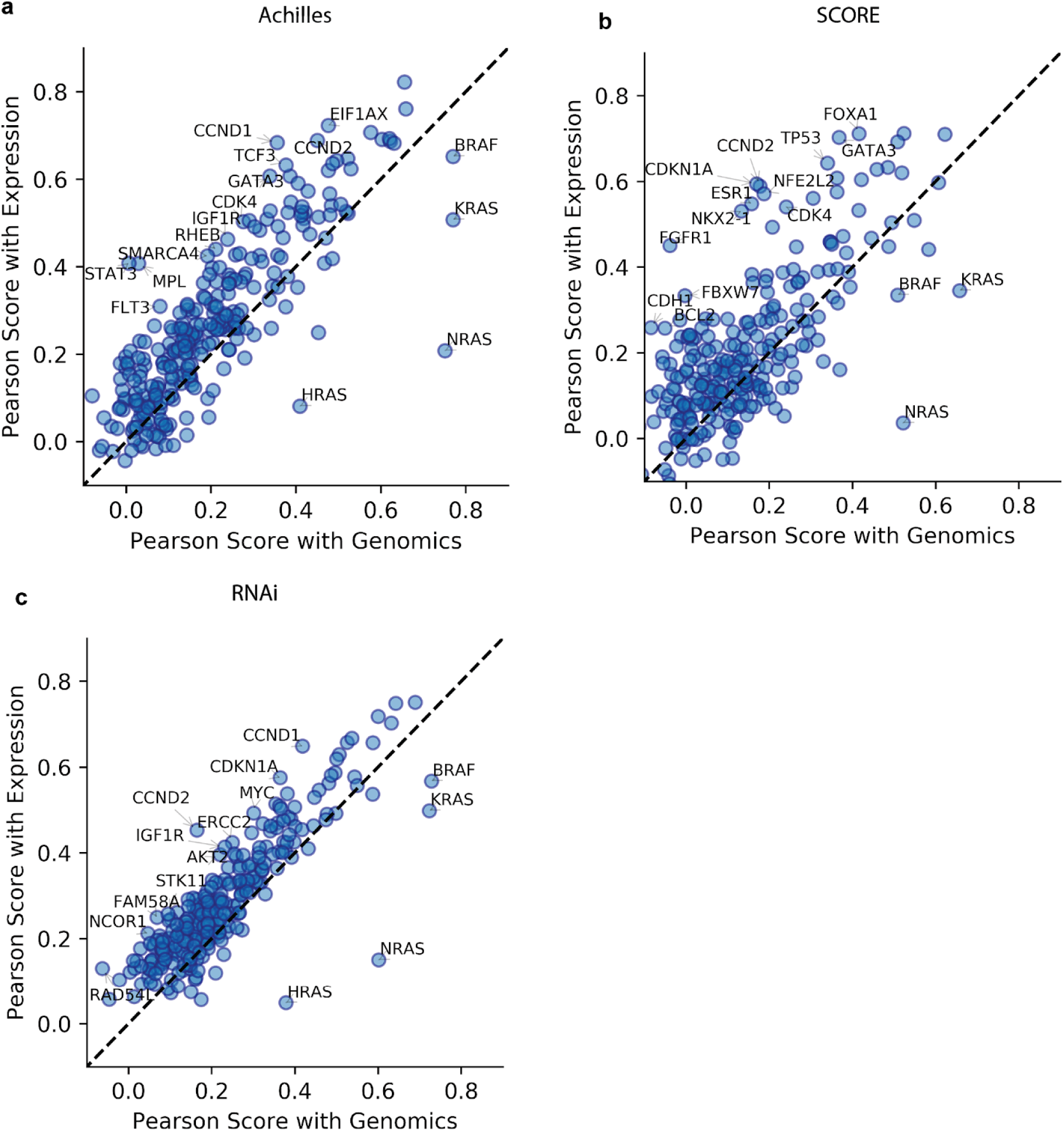
Predictive performance of dependency on OncoKB oncogenes. **a.** Comparison of predictive performance using either DNA (x-axis) or expression (y-axis) features in Achilles. **b**. Similar for Score. **c**. Similar for RNAi.

**Fig. 2 - figure supplement 3:**
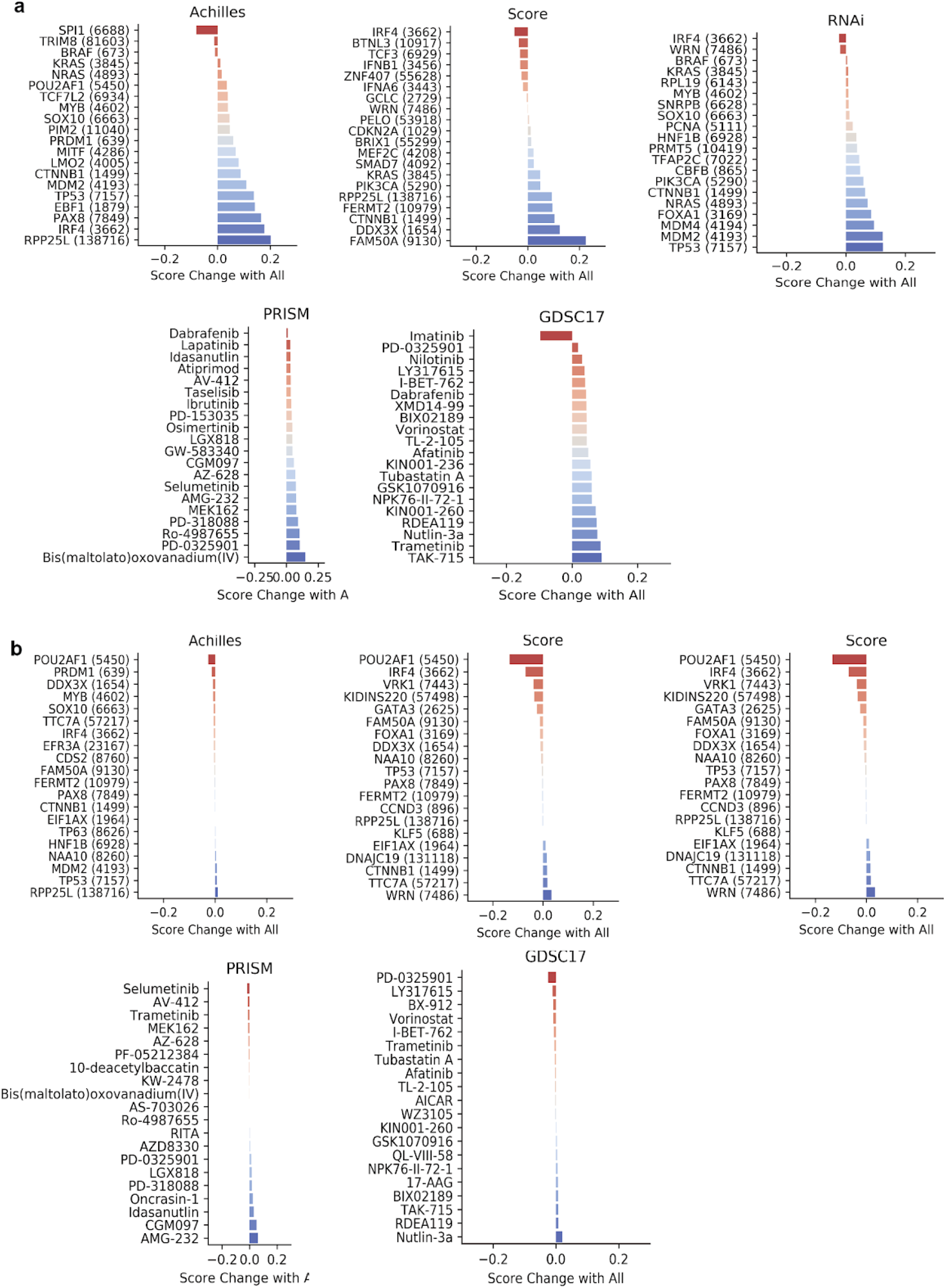
Benefits of including expression or DNA features. **a.** For the perturbations best predicted with DNA features, the change in model performance when expression features are added. **b**. The reverse.

**Fig. 4 - figure supplement 1:**
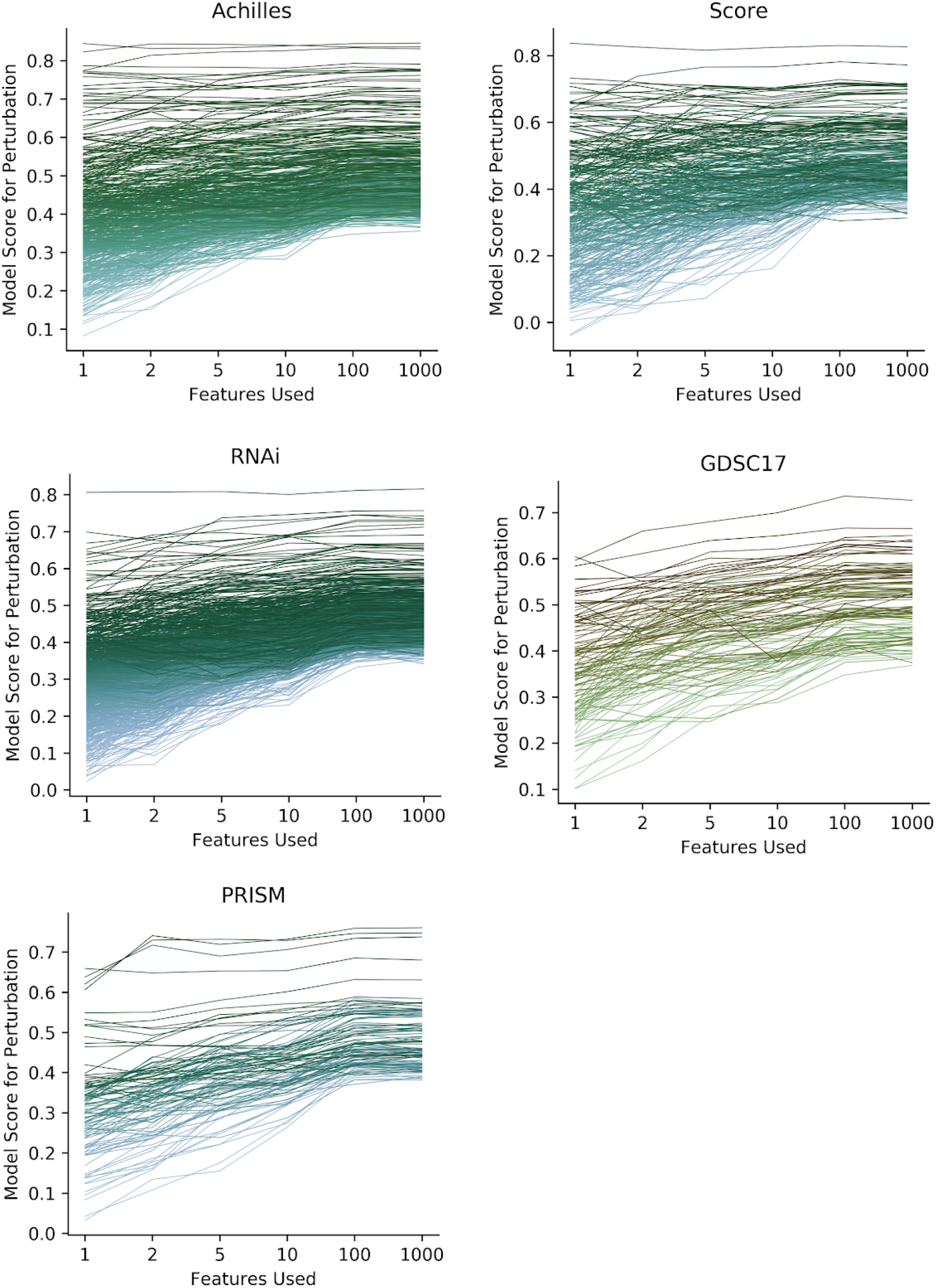
Model performance with variable numbers of features provided. Each line represents the performance of models using all feature types for a specific perturbation. Perturbations were restricted to those originally predicted with Pearson correlation greater than 0.4. For each cross-validation fold, the 1000 features with highest correlation to the target variable were first selected and a random forest trained. Features were ranked by importance in the trained model and additional models trained with only the indicated subset of features.

**Fig. 4 - figure supplement 2:**
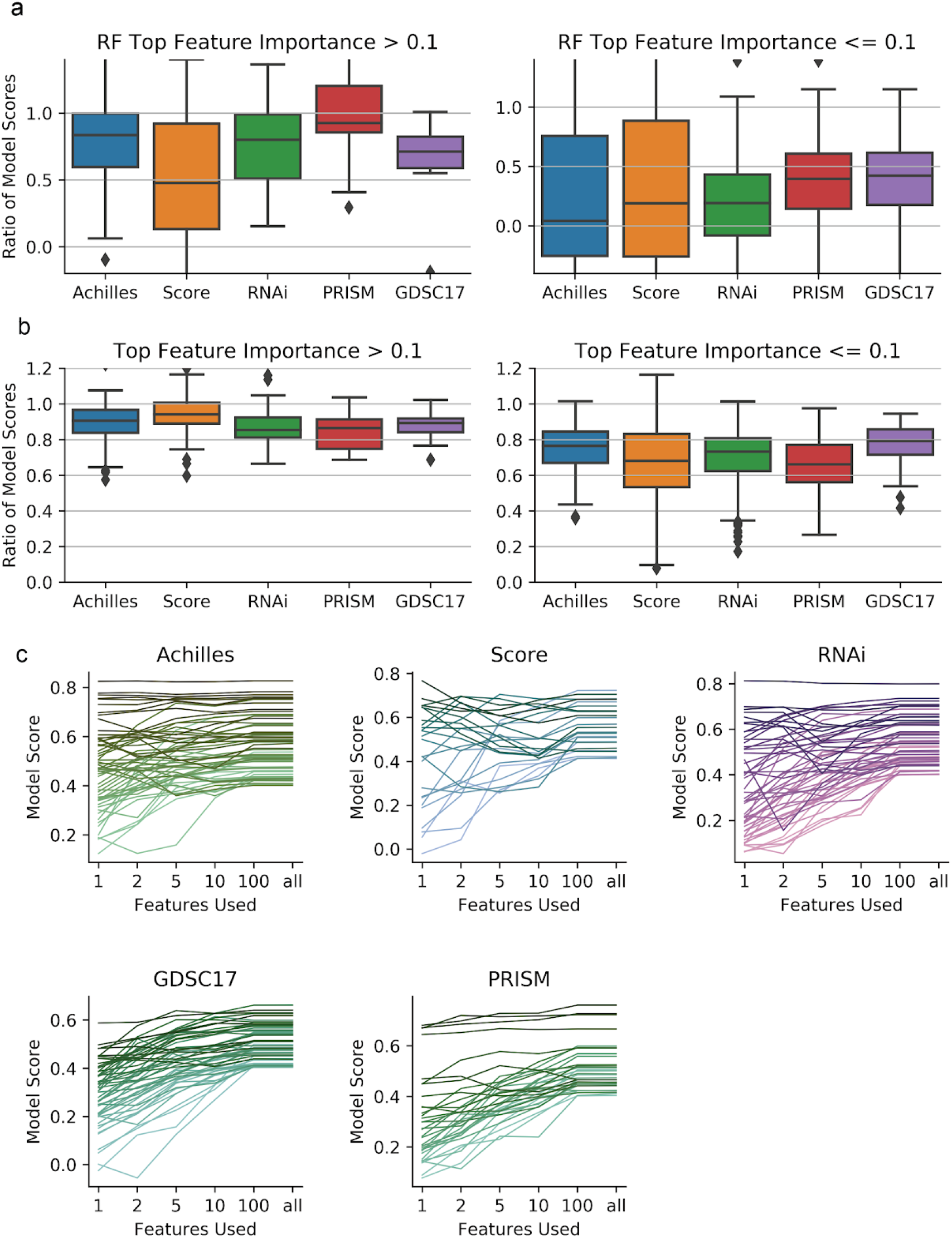
Additional ablation performance measures. **a.** The ratio of random forest predictor scores using all features or using only two features selected using random forest feature importance. Only predictors scoring above 0.4 are included. **b.** Similar, using elastic net predictors with one or all features. **c.** Score of the elastic net as a function of number of features used. Each line represents the performance of a predictor using all feature types. For each cross-validation fold using all features, features were then ranked by normalized coefficient in the trained model and additional models trained with only the indicated subset of features.

**Fig. 6 - figure supplement 1:**
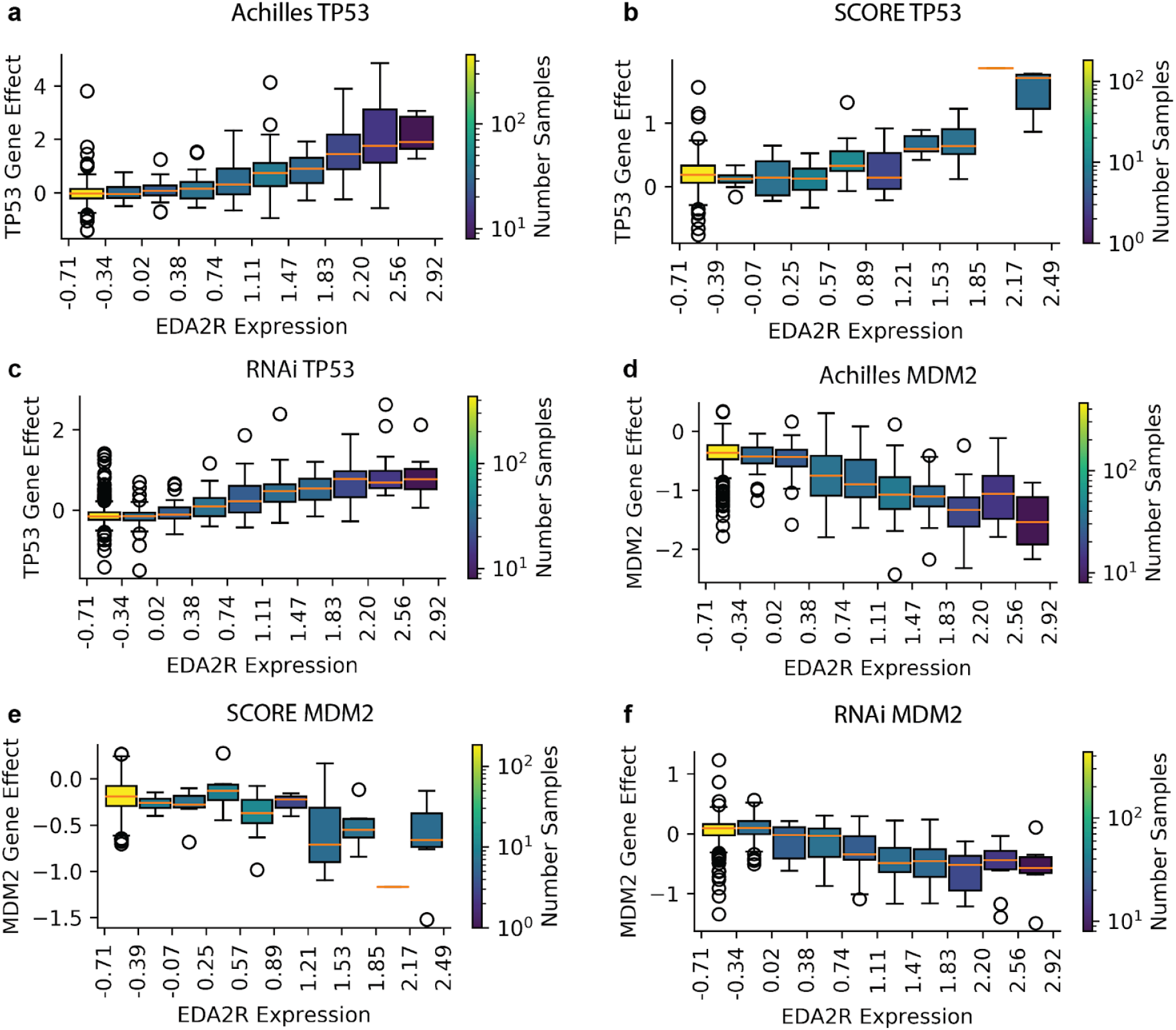
Relationship of EDA2R expression with post-perturbation viability of TP53 and MDM2 in genetic datasets.

**Fig. 6 - figure supplement 2:**
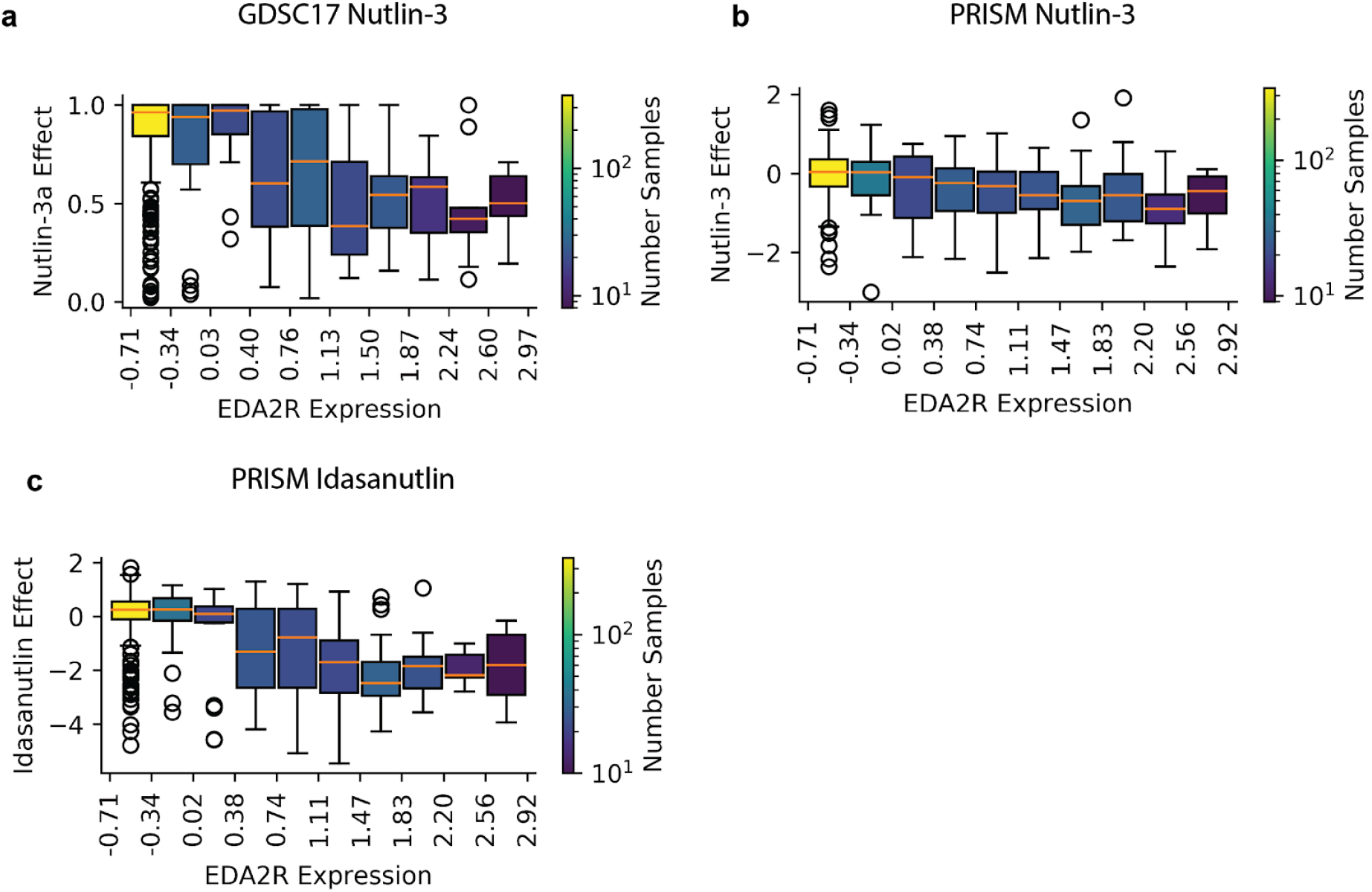
Relationship of EDA2R expression with post-perturbation viability after MDM2 inhibitors.

**Fig. 6 - figure supplement 3:**
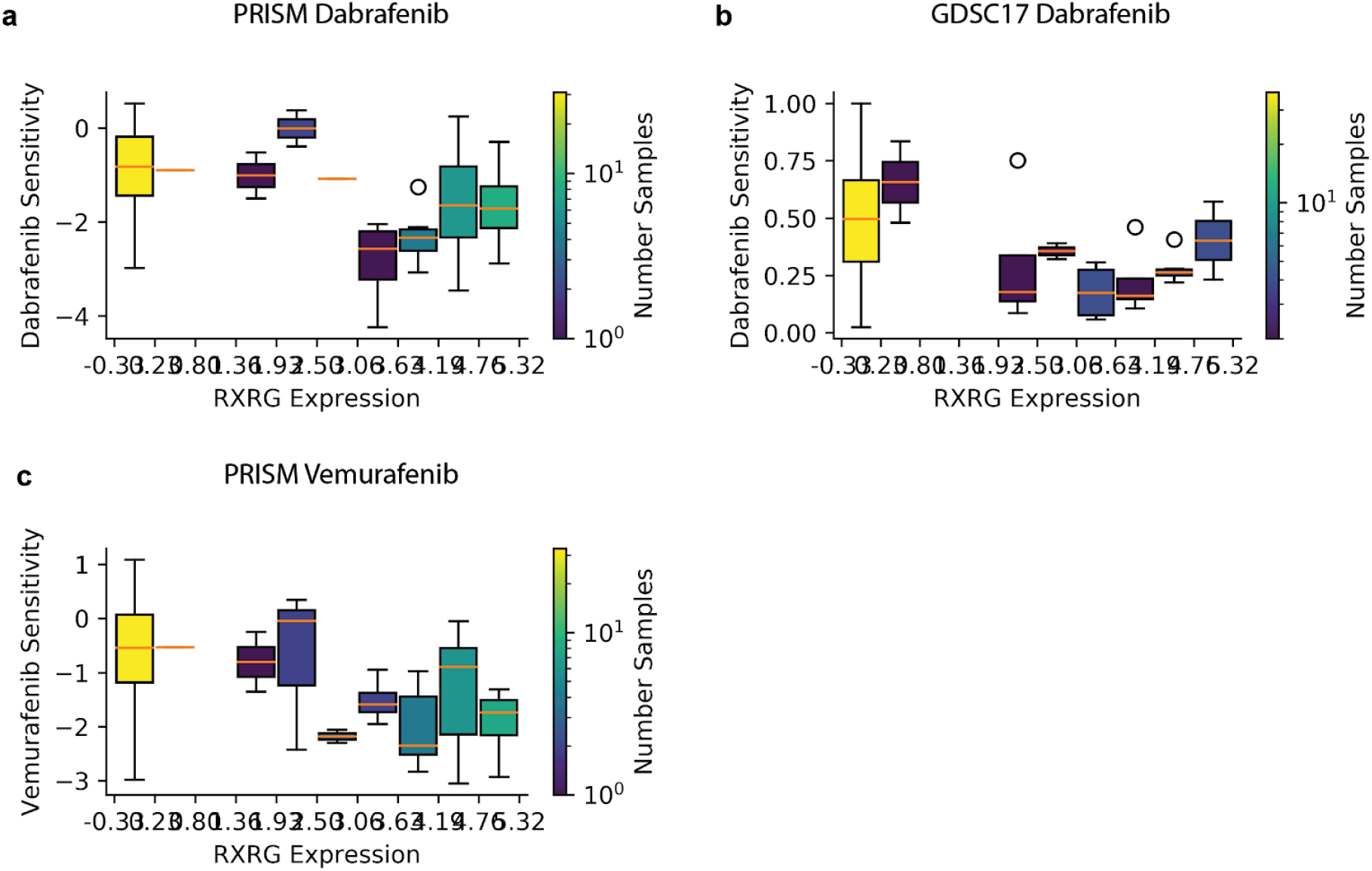
Relationship of viability after treatment with BRAF inhibitors with RXRG expression in the cell lines with hotspot BRAF mutations.

**Fig. 6 - figure supplement 4:**
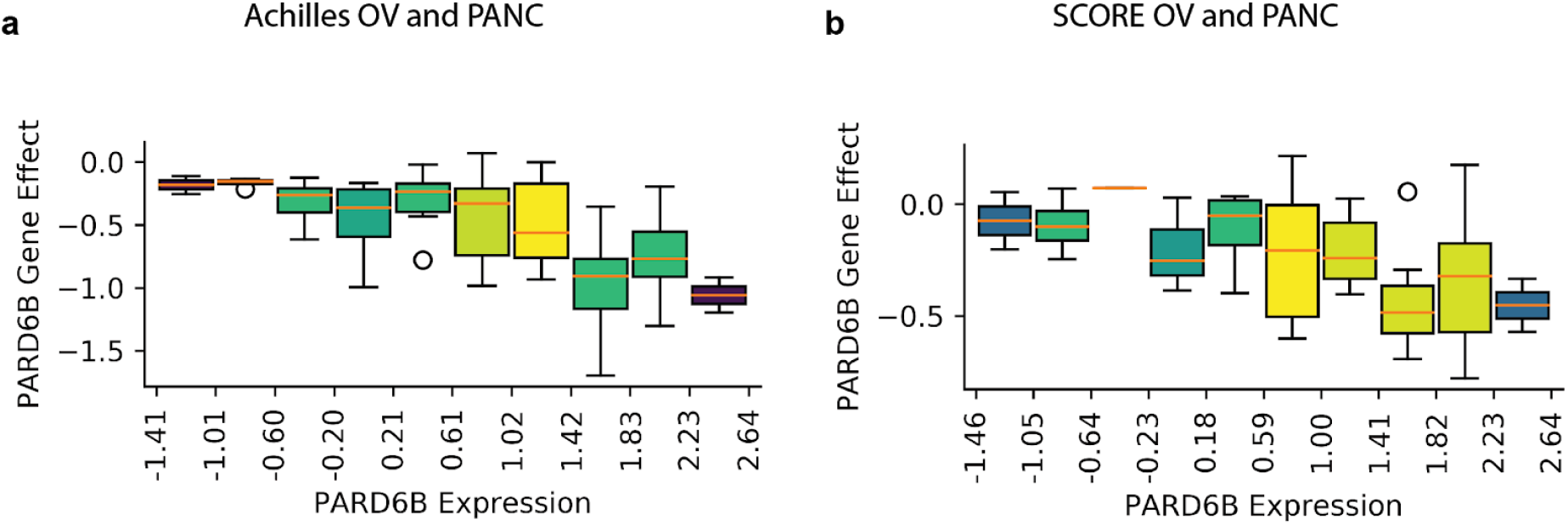
Relationship of viability after knockout of PARD6B with PARD6B expression in pancreatic and ovarian cancer cell lines.

## Notes

### Summary of Updates

Reformatting.

## References

1. Macconaill LE, Garraway LA. Clinical implications of the cancer genome. J Clin Oncol. 2010;28: 5219–5228.

2. Haber DA, Gray NS, Baselga J. The evolving war on cancer. Cell. 2011;145: 19–24.

3. Cheng ML, Berger MF, Hyman DM, Solit DB. Clinical tumour sequencing for precision oncology: time for a universal strategy. Nat Rev Cancer. 2018;18: 527–528.

4. Prasad V. Perspective: The precision-oncology illusion. Nature. 2016;537: S63.

5. Letai A. Functional precision cancer medicine—moving beyond pure genomics. Nat Med. 2017;23: 1028–1035.

6. Zhu B, Song N, Shen R, Arora A, Machiela MJ, Song L, et al. Integrating Clinical and Multiple Omics Data for Prognostic Assessment across Human Cancers. Sci Rep. 2017;7: 16954.

7. Ali M, Aittokallio T. Machine learning and feature selection for drug response prediction in precision oncology applications. Biophys Rev. 2019;11: 31–39.

8. Ding Z, Zu S, Gu J. Evaluating the molecule-based prediction of clinical drug responses in cancer. Bioinformatics. 2016;32: 2891–2895.

9. Costello JC, NCI DREAM Community, Heiser LM, Georgii E, Gönen M, Menden MP, et al. A community effort to assess and improve drug sensitivity prediction algorithms. Nature Biotechnology. 2014. pp. 1202–1212. doi:10.1038/nbt.2877

10. Franco M, Jeggari A, Peuget S, Böttger F, Selivanova G, Alexeyenko A. Prediction of response to anti-cancer drugs becomes robust via network integration of molecular data. Sci Rep. 2019;9: 2379.

11. Wang X, Sun Z, Zimmermann MT, Bugrim A, Kocher J-P. Predict drug sensitivity of cancer cells with pathway activity inference. BMC Med Genomics. 2019;12: 15.

12. Ben-Hamo R, Jacob Berger A, Gavert N, Miller M, Pines G, Oren R, et al. Predicting and affecting response to cancer therapy based on pathway-level biomarkers. Nat Commun. 2020;11: 3296.

13. Iorio F, Knijnenburg TA, Vis DJ, Bignell GR, Menden MP, Schubert M, et al. A Landscape of Pharmacogenomic Interactions in Cancer. Cell. 2016;166: 740–754.

14. Aben N, Vis DJ, Michaut M, Wessels LFA. TANDEM: a two-stage approach to maximize interpretability of drug response models based on multiple molecular data types. Bioinformatics. 2016;32: i413–i420.

15. Geeleher P, Cox NJ, Huang RS. Clinical drug response can be predicted using baseline gene expression levels and in vitro drug sensitivity in cell lines. Genome Biol. 2014;15: R47.

16. Rodon J, Soria J-C, Berger R, Miller WH, Rubin E, Kugel A, et al. Genomic and transcriptomic profiling expands precision cancer medicine: the WINTHER trial. Nat Med. 2019;25: 751–758.

17. Behan FM, Iorio F, Picco G, Gonçalves E, Beaver CM, Migliardi G, et al. Prioritization of cancer therapeutic targets using CRISPR-Cas9 screens. Nature. 2019;568: 511–516.

18. Corsello SM, Nagari RT, Spangler RD, Rossen J, Kocak M, Bryan JG, et al. Non-oncology drugs are a source of previously unappreciated anti-cancer activity. Cancer Biology. bioRxiv; 2019. p. 589.

19. McFarland JM, Ho ZV, Kugener G, Dempster JM, Montgomery PG, Bryan JG, et al. Improved estimation of cancer dependencies from large-scale RNAi screens using model-based normalization and data integration. Nat Commun. 2018;9: 4610.

20. Li Y, Umbach DM, Krahn J, Shats I, Li X, Li L. Predicting Tumor Response to Drugs based on Gene-Expression Biomarkers of Sensitivity Learned from Cancer Cell Lines. 2020. p. 2020.07.03.180620. doi:10.1101/2020.07.03.180620

21. Liberzon A, Subramanian A, Pinchback R, Thorvaldsdóttir H, Tamayo P, Mesirov JP. Molecular signatures database (MSigDB) 3.0. Bioinformatics. 2011;27: 1739–1740.

22. Chakravarty D, Gao J, Phillips SM, Kundra R, Zhang H, Wang J, et al. OncoKB: A Precision Oncology Knowledge Base. JCO Precis Oncol. 2017;2017. doi:10.1200/PO.17.00011

23. Zerbino DR, Achuthan P, Akanni W, Amode MR, Barrell D, Bhai J, et al. Ensembl 2018. Nucleic Acids Res. 2018;46: D754–D761.

24. Li T, Wernersson R, Hansen RB, Horn H, Mercer J, Slodkowicz G, et al. A scored human protein-protein interaction network to catalyze genomic interpretation. Nat Methods. 2017;14: 61–64.

25. Giurgiu M, Reinhard J, Brauner B, Dunger-Kaltenbach I, Fobo G, Frishman G, et al. CORUM: the comprehensive resource of mammalian protein complexes—2019. Nucleic Acids Res. 2018;47: D559–D563.

26. Klebanov L, Yakovlev A. Diverse correlation structures in gene expression data and their utility in improving statistical inference. Ann Appl Stat. 2007;1: 538–559.

27. Zoppoli G, Regairaz M, Leo E, Reinhold WC, Varma S, Ballestrero A, et al. Putative DNA/RNA helicase Schlafen-11 (SLFN11) sensitizes cancer cells to DNA-damaging agents. Proc Natl Acad Sci U S A. 2012;109: 15030–15035.

28. Warters RL, Packard AT, Kramer GF, Gaffney DK, Moos PJ. Differential gene expression in primary human skin keratinocytes and fibroblasts in response to ionizing radiation. Radiat Res. 2009;172: 82–95.

29. Macaeva E, Saeys Y, Tabury K, Janssen A, Michaux A, Benotmane MA, et al. Radiation-induced alternative transcription and splicing events and their applicability to practical biodosimetry. Sci Rep. 2016;6: 19251.

30. Giacomelli AO, Yang X, Lintner RE, McFarland JM, Duby M, Kim J, et al. Mutational processes shape the landscape of TP53 mutations in human cancer. Nat Genet. 2018;50: 1381–1387.

31. Chène P. Inhibiting the p53–MDM2 interaction: an important target for cancer therapy. Nat Rev Cancer. 2003;3: 102–109.

32. Rambow F, Rogiers A, Marin-Bejar O, Aibar S, Femel J, Dewaele M, et al. Toward Minimal Residual Disease-Directed Therapy in Melanoma. Cell. 2018;174: 843–855.e19.

33. Rambow F, Marine J-C, Goding CR. Melanoma plasticity and phenotypic diversity: therapeutic barriers and opportunities. Genes Dev. 2019;33: 1295–1318.

34. Rydenfelt M, Wongchenko M, Klinger B, Yan Y, Blüthgen N. The cancer cell proteome and transcriptome predicts sensitivity to targeted and cytotoxic drugs. Life Sci Alliance. 2019;2. doi:10.26508/lsa.201900445

35. Paull EO, Aytes A, Subramaniam P, Giorgi FM. A Modular Master Regulator Landscape Determines the Impact of Genetic Alterations on the Transcriptional Identity of Cancer Cells. bioRxiv. 2019. Available: https://www.biorxiv.org/content/10.1101/758268v1.abstract

36. Alvarez MJ, Shen Y, Giorgi FM, Lachmann A, Ding BB, Ye BH, et al. Functional characterization of somatic mutations in cancer using network-based inference of protein activity. Nat Genet. 2016;48: 838–847.

37. DepMap B. DEMETER 2 Combined RNAi. 2019. doi:10.6084/m9.figshare.9170975.v1

38. CancerRxGene. CancerRxGene/gdscIC50. In: GitHub [Internet]. [cited 25 Oct 2018]. Available: https://github.com/CancerRxGene/gdscIC50

39. DepMap B. Project SCORE processed with CERES. 2019. doi:10.6084/m9.figshare.9116732.v1

40. DepMap B. DepMap 19Q2 Public. 2019. doi:10.6084/m9.figshare.8061398.v1

41. Dempster JM, Rossen J, Kazachkova M, Pan J, Kugener G, Root DE, et al. Extracting Biological Insights from the Project Achilles Genome-Scale CRISPR Screens in Cancer Cell Lines. bioRxiv. 2019. p. 720243. doi:10.1101/720243

42. McDonald ER 3rd, de Weck A, Schlabach MR, Billy E, Mavrakis KJ, Hoffman GR, et al. Project DRIVE: A Compendium of Cancer Dependencies and Synthetic Lethal Relationships Uncovered by Large-Scale, Deep RNAi Screening. Cell. 2017;170: 577–592.e10.

